# The independent and combined influence of schizophrenia polygenic risk score and heavy cannabis use on risk for psychotic disorder: A case-control analysis from the EUGEI study

**DOI:** 10.1101/844803

**Authors:** Marta Di Forti, Beatrice Wu-Choi, Diego Quattrone, Alexander L Richards, Tom P Freeman, Giada Tripoli, Charlotte Gayer-Anderson, Victoria Rodriguez, Hannah Jongsma, Laura Ferraro, Caterina La Cascia, Sarah Tosato, Ilaria Tarricone, Domenico Berardi, Andrei Szoke, Celso Arango, Julio Bobes, Julio Sanjuan, Jose Luis Santos, Manuel Arrojo, Eva Velthorst, Miquel Bernardo, Cristina Marta Del Ben, Jean-Paul Selten, Peter B Jones, James B Kirkbride, Bart NP Rutten, Lieuwe de Haan, Jim van os, Michael Lynskey, Craig Morgan, Evangelos Vassos, Michael O’Donovan, Cathryn Lewis, Pak C Sham, Robin M Murray, EU-GEI WP2 Group

## Abstract

**Background:** Some recent studies have challenged the direction of causality for the association between cannabis use and psychotic disorder, suggesting that cannabis use initiation is explained by common genetic variants associated with risk of schizophrenia. We used data from the European Union Gene-Environment Interaction consortium (EUGEI) case-control study to test for the independent and combined effect of heavy cannabis use, and of Schizophrenia Polygenic risk score (SZ PRS), on risk for psychotic disorder.

**Methods:** Genome-wide data were obtained from 492 first episode psychosis patients (FEPp) and from 787 controls of European Ancestry, and used to generate SZ PRS from the summary results of an independent meta-analysis. Information on pattern of cannabis use was used to build a 7-level *frequency-type* composite cannabis use measure that we previously found was a strong predictor of psychotic disorder.

**Results:** SZ PRS did not predict cannabis initiation (b=0.027; p=0.51) or how frequently controls (b=0.027; p=0.06) or FEPp (b=0.006; p=0.91) used it, or the type of cannabis they used (Controls: b = 0.032; p=0.31); FEPp: b= 0.005; p=0.89). The *frequency-type* composite cannabis use measure (OR=1.32; 95% CI 1.22-1.44) and SZ PRS (OR=2.29; 95%CI 1.71-3.05) showed independent effects from each other on the OR for psychotic disorder.

**Conclusion:** SZ PRS does not predict an individual’s propensity to try cannabis, frequency of use, or the potency of the cannabis used. Our findings provide the first evidence that SZ PRS and heavy cannabis use exert effects independent from each other on the risk for psychotic disorder.

**Funding source:** Medical Research Council (MRC). The European Community’s Seventh Framework Program grant (agreement No. HEALTH-F2-2009-241909 [Project EU-GEI]). São Paulo Research Foundation under grant number 2012/0417-0. National Institute for Health Research (NIHR) Biomedical Research Centre (BRC) at South London and Maudsley NHS Foundation Trust and King’s College London and the NIHR BRC at University College London. Wellcome Trust (grant 101272/Z/12/Z).

## Introduction

Cannabis is used by some 200 million people worldwide, and its use and potency have increased in many countries ^1-3^. Prospective epidemiological studies ^4^, as well as biological investigations ^5^, demonstrate a causal link between cannabis use and psychotic disorder. Recent evidence has confirmed a) a dose–response association with the highest odds of psychotic disorder in the heaviest cannabis users ^6^ and b) that high potency cannabis carries the greatest risk for psychotic disorder ^7^. Indeed, daily cannabis use and use of high potency types have been linked to variation in the incidence of psychotic disorder across Europe ^6^.

A recent study ^8^ showed that individuals with a family history of schizophrenia who develop a cannabis induced psychotic disorder, are especially likely to transition to schizophrenia. However, not all heavy cannabis users develop a psychotic disorder in the first place, and it remains unclear which genetic factors influence individual vulnerability to the psychotogenic effects of cannabis use.

Patterns of cannabis use such as lifetime cannabis use (never/ever used) and Cannabis Use Disorder (CUD) are influenced by genetic factors ^9,10^. Twin heritability reaches 45% for lifetime cannabis use and 51% to 70% for CUD ^11,12^. Genome wide Association Studies (GWAS) have also shown a significant genetic correlation between lifetime cannabis use and CUD on the one hand, and schizophrenia on the other ^13^. Moreover, polygenic risk scores for schizophrenia (SZ PRS) have been reported to explain a small but significant proportion of the variance in lifetime cannabis use, quantity of cannabis used ^14^ and CUD ^13^.

Mendelian Randomization (MR) studies have investigated if the reported genetic association between cannabis use phenotypes and schizophrenia results from a causal relationship between the two or from genetic pleiotropy; findings have been contradictory ^15-17^. In the most recent study, Pasman et al ^17^, used data from a large GWAS of cannabis use initiation to perform a bi-directional two sample MR analysis. In contrast to Vaucher et al ^15^, they suggest a causal positive association of schizophrenia genes on cannabis initiation but not vice versa.

However, so far MR studies have only been able to explore a causal association between schizophrenia genes and cannabis use initiation rather than with those patterns of heavy cannabis use shown to impact on risk of psychotic disorder.

Therefore, using detailed data on pattern of cannabis use and GWAS data from a large multisite study, we aimed to test: 1) if SZ PRS predict cannabis initiation and/or patterns of cannabis use in population controls and first episode psychosis patients; 2) the individual and combined effects of SZ PRS and cannabis use on the risk of psychotic disorder and 3) if adding SZ PRS data to information on patterns of cannabis use improves the identification of those heavy cannabis users who will develop psychotic disorder.

## Methods and materials

This paper derives from analyses of the EUGEI first episode case-control samples recruited between 1/5/2010 and 1/4/2015 in 17 catchment areas in England, France, the Netherlands, Italy, Spain and Brazil^18^.

### Participants

#### Cases

Patients presenting with their first episode of psychosis (FEPp) were identified by trained researchers who carried out regular checks across the 17 catchment area Mental Health Services. FEPp were included if a) age 18-64 years and b) resident within the study areas at the time of their first presentation, and received a diagnosis of psychosis (ICD-10 F20-33); further details are provided in the supplementary methods and in our recent publication ^7^. Using the Operational Criteria Checklist algorithm ^19,20^, all cases interviewed received a research-based diagnosis. FEPs were excluded if a) previously treated for psychosis, b) they met criteria for organic psychosis (ICD-10: F09), or for a diagnosis of transient psychotic symptoms resulting from acute intoxication (ICD-10: F1X.5).

#### Controls

Random and Quota sampling strategies were adopted to guide the recruitment of controls from each of the sites. The most accurate local demographic data available were used to set quotas for controls to ensure the samples’ representativeness of each catchment area’s population at risk (see supplementary material). Controls were excluded if they had received a diagnosis of, and/or treatment for, psychotic disorder.

All participants provided informed, written consent. Ethical approval was provided by relevant research ethics committees in each of the study sites.

#### Measures of cannabis use

Data on patterns of cannabis use were collected using the modified Cannabis Experience Questionnaire further updated (CEQ_EU-GEI_)^7^. None of the materials we used for the participants recruitment referred to cannabis or to its potential role as a risk factor for psychotic disorder. Participants were asked if they had ever used cannabis. If yes, they were asked to answer questions about their pattern of use, including the type of cannabis allowing the participants to report the “street” name, in the original language, of the cannabis they used with no reference to its potency.

We used measures of cannabis use that, in an independent sample, we reported ^21^ to increase the ORs for Psychotic Disorder: I) Age at first use of cannabis; II) lifetime frequency of use and III) the potency of the cannabis used. The latter was estimated, as described in Di Forti et al ^7^ using the EMCDDA 2016 report ^22^ and additional National published data on the concentration (%) of Tetrahydrocannabinol (THC) expected in the different types of cannabis available across Europe ^22,23-30^ (see supplementary material). We also kept the variable “lifetime” ever cannabis use Yes/No to be able to compare our findings to the existing literature on the genetics of cannabis initiation ^17^. Finally, we used the lifetime frequency of use and the cannabis potency variables to build the ““*frequency-type* composite cannabis use measure” that we previously tested ^21^ and replicated ^5^ to be a strong predictor of psychotic disorder. The “*frequency-type”* composite cannabis use measure includes 7 scores associated with a steady (from 0 to 6) increase in the OR for psychotic disorder ^7,21^: never used cannabis= 0; rare use of low potency cannabis (THC<10%)=1; rare use of high potency cannabis (THC=>10%)=2; use>than once a week of low potency cannabis (THC<10%)=3; use>than once a week of low potency cannabis (THC=>10%)=4; daily use of low potency cannabis (THC<10%)=5; daily use of high potency cannabis (THC=>10%)=6.

#### Genotyping

Samples were genotyped at the MRC Centre for Neuropsychiatric Genetics and Genomics in Cardiff (UK) using a custom Illumina HumanCoreExome-24 BeadChip genotyping array covering 570,038 genetic variants. To identify ethnic groups, we combined our dataset with the 1000 Genome Project (1000G), phase 3 and performed Principal Component Analysis on the overlapping SNPs. We then used the first two principal components to carry out 4-means clustering which identified the main study ethnic ancestry groups: African, European and Asian (**Figure 1**). For example, individuals of European ancestry were defined as having PC values within 6 standard deviations from the mean PC of the EUR in 1000G, and retained for the downstream analyses.

**Figure 1:**
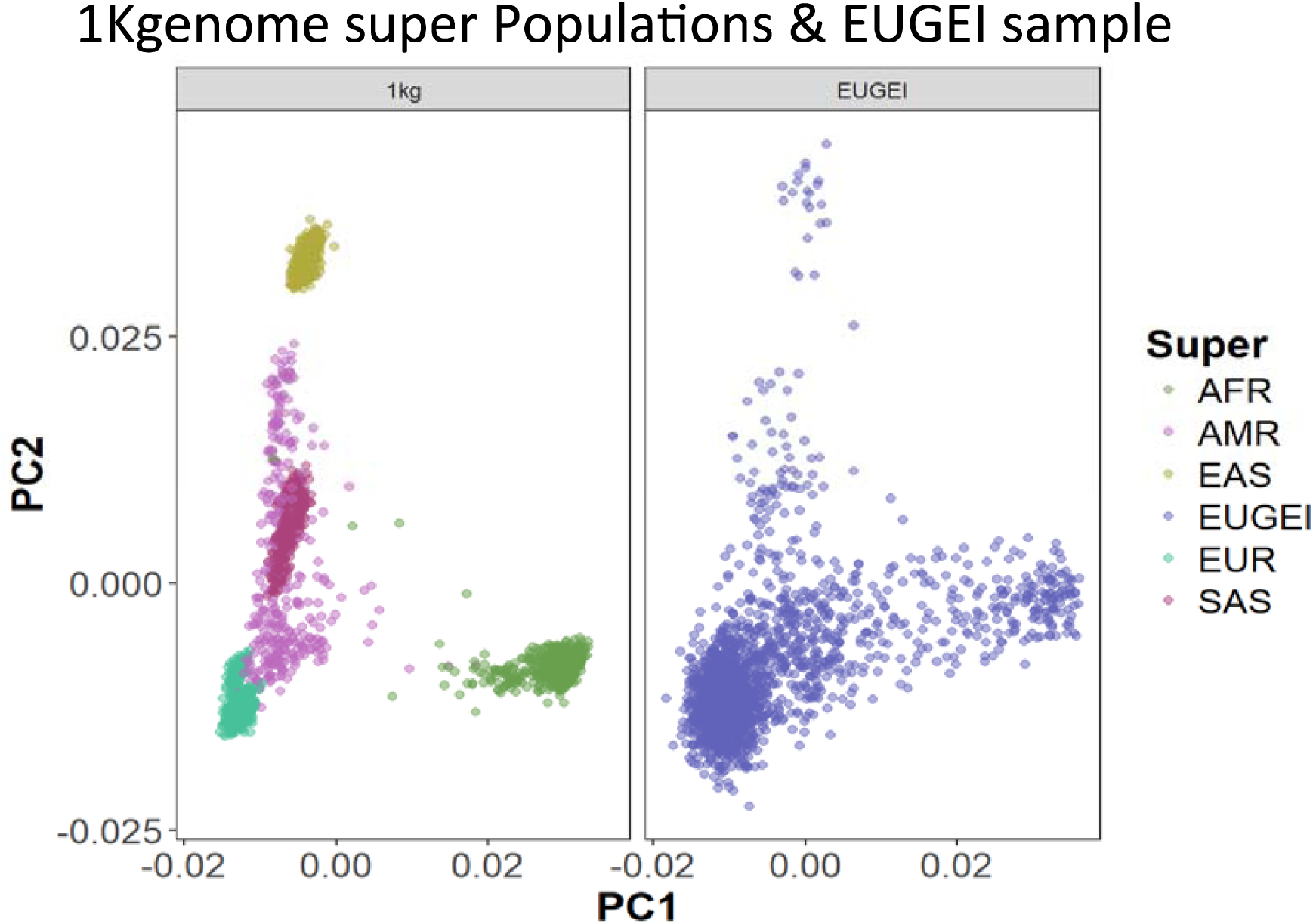
This plot shows how the first two Principal Component (PC1 and PC2) separate the 1Kgenome project 5 super-populations (AFR: African; AMR: Ad Mixed American; EAS: East Asian; EUR: European; SAS: South Asian) and the main populations of the EUGEI (European Union Gene-Environment Interaction consortium) sample.

#### Statistical Analysis

Polygenic risk scores (PRS) for schizophrenia (SZ PRS) were generated using PRSice from the summary results of the PGC analysis of schizophrenia, wave 2 ^31^. Clumping was performed to obtain SNPs in approximate linkage disequilibrium with an r^2^ < 0.25 within a 250 kb window. PRS were calculated, separately for each of the three ancestry populations, at P-value thresholds of 0.05 ^31^. Then, each PRS was standardised (std_PRS) to mean of 0 and standard deviation of 1, excluding the MHC region. ^32^. In STATA 15 we also calculated SZ PRS *quintiles.*

Adjusted logistic regression models were run to estimate: 1) if SCZ PRS predicted life time cannabis use and/or pattern of cannabis use and 2) the independent and combined effect of the selected measures of cannabis use, and the SZ PRS on the ORs for psychotic disorder. We fitted interaction terms to the logistic models and used likelihood ratio tests, to test if SZ PRS modify the effect of 1) the individual measures of cannabis use and 2) the “*frequency-type* composite cannabis use measure on the ORs for psychotic disorder. All regression models were adjusted for: 10 PCs, sites, age, sex and tobacco smoking as defined in our previous publication ^7^ (0=never smoked or <=10 cigarettes x day; 1= 11 cigarettes or more x day). In STATA 15 we used the “*marginplot*” command to display graphically the average predicted probability (y=axis) of being a case over increasing values of SZ PRS (x=axis) across different levels of exposures to cannabis use (i.e. *frequency-type* composite cannabis use measure).

Finally, the STATA 15 *lroc* command was used to assess the discriminatory ability (correctly classify *case_control* status) of some of the models tested.

## Results

We approached 1519 patients FEP patients; 356 (23%) refused to participate, 19 (1%) could not consent because of language barriers and 14 (0.9%) were excluded as they did not meet the age inclusion criteria. Patients who refused to participate were older, more likely to be women and of European ancestry (supplementary methods). 1130 FEPp and 1499 population controls consented to take part. DNA samples were successfully collected from N=2190 participants out of the total N=2629 recruited (83%); DNA was extracted from blood (N=1857) or saliva (N=312).

The GWAS call rate of 98% (N=2125) resulted in a total sample with available genetic data to build the SZ PRS of FEPp=999 and controls=1147. Using the PCs approach described in the methods we calculated the Nagelkerke R^2^ by the SZ PRS in each of our main Ethnic Ancestry groups: African (R^2^=0.03%; p=0.437; Controls N=301; FEPp N=402), European (R^2^=6.3%; p=5.15E-14; Controls N=787; FEPp N=492) and Others (R^2^=5.2%; p=1.03E-08; Controls N=59; FEPp N=105).

These differences in R^2^ across the 3 main ethnic groups reflect the over-representation of individuals of European Ancestry in the PGC2 training sample used to calculate the SZ PRS and are consistent with previous reports ^33^. Therefore, we restricted the working sample to those of white European Ancestry (supplementary methods flow chart). As shown in **Figure 2** the SZ PRS was on average higher in FEPp than in controls (FEPp mean SZ PRS=0.254, SD 0.97; controls mean SZ PRS= −0.16, SD 0.98; t= −7.37,df=1477; p<0.001]. Consistently, there were more controls in the SZ PRS *quintile1* compared to FEPp [Controls: 190/256(24.1%); FEPp: 66/256(13.4%); p<0.001]; in contrast there was a larger proportion of FEPp compared to controls in *quintile5* [FEPp: 139/256 (28.25%); Controls: 116/256(14.74%); p<0.001].

**Figure 2.**
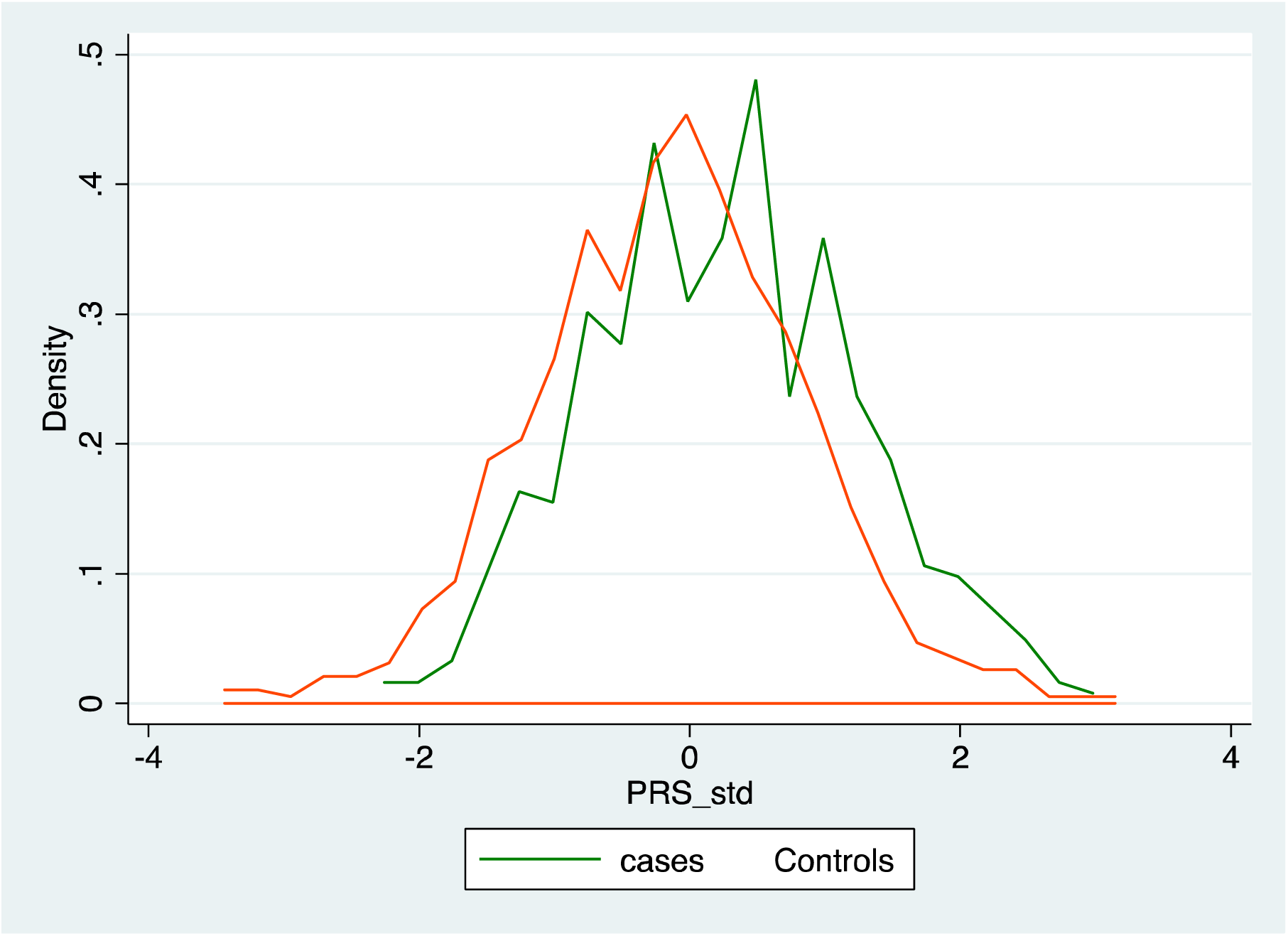
SZ PRS (PRS_std) distribution in FEP (green line) and controls (red line) in the sub-sample sample of European Ancestry

The final sample consisted of 492 first episode psychosis patients (FEPp) and 787 controls. FEPp were younger (FEPp mean age =32.3, SD 11.0; controls mean age= 37.5, SD 13.1; p<0.001) and more likely to be male than controls [FEPp: 300(60.9%); controls: 373 (47.4%); p<0.001]. FEPp were also more likely to have ever tried cannabis [FEPp: 332 (67.5%); Controls: 418 (53.1%); p<0.001], to have first used it at age 15 years old or younger [FEPp: 135(27.4%); controls: 101(12.8%); p<0.001] and to have used it daily [FEPp: 148 (30.0%); controls: 46 (5.9%); p<0.001]. FEPp were also more likely to have tried more potent types [FEPp: 223 (45.2%); controls: 185(23.5%); p<0.001] and to have used potent types daily [FEPp: 117 (20.7%); controls: 28(3.5%); p<0.001] than controls (**Table 1**). FEPp were also more likely to have smoked 11 or more tobacco cigarettes daily compared to controls [FEPp: 328(66.8%); controls: 266 (33.8%); p<0.001].

**Table 1:** Differences between first episode psychosis patients (FEPp) and controls in basic socio-demographics and patterns of cannabis use

|  | **Controls**<br>**N= 787** | **FEPp**<br>**N=492** | **df** | **Test statistics** | **P value** |
| --- | --- | --- | --- | --- | --- |
| **Age (mean; sd)** | 37.5 (13.1) | 32.3 (11.0) | 1477 | t = 7.43 | <0.001 |
| **Gender (male ;N)** | 47.4 (373) | 60.9(300) | 1 | Chi2 = 22.39 | <0.001 |
| **Life time cannabis use (%; N)**<br>**Yes** | 53.1 (418) | 67.5(332) | 1 | Chi2 = 25.10 | <0.001 |
| **Age at 1<sup>st</sup> use =< 15 years old**<br>**(% of the total; N)** | 12.8 (101) | 27.4 (135) | 2 | Chi2 = 53.11 | <0.001 |
| **Frequency of use (%; N)**<br>**Never used** | 46.9 (369) | 32.5 (160) |  |  |  |
| **Less than weekly (rare use)** | 40.0 (314) | 25.4 (124) | 3 | Chi2=146.64 | <0.001 |
| **More than once x week** | 7.2 (58) | 12.1 (60) |  |  |  |
| **Daily** | 5.9 (46) | 30.0 (148) |  |  |  |
| **Potency of cannabis used (%;N)**<br>**Never Used** | 46.9(369) | 32.5(160) |  |  |  |
| **THC<10%** | 29.6(233) | 22.3 (109) | 2 | Chi2= 43.80 | <0.001 |
| **THC=>10%** | 23.5 (185) | 45.2(223) |  |  |  |
| **Type-Frequency composite cannabis use measure**<br>**Never used** | 46.9(369) | 32.5(160) |  |  |  |
| **Rare Use of THC<10%** | 22.5 (175) | 10.0(49) |  |  |  |
| **Rare Use of THC=>10%** | 16.6(131) | 13.5(66) |  |  |  |
| **Use > than once x week of THC<10%** | 4.6(37) | 5.0(25) | 6 | Chi2= 137.16 | <0.001 |
| **Use > than once x week of THC=>10%** | **3.2(25)** | **8.0(39)** |  |  |  |
| **Daily use of THC<10%** | **2.8(22)** | **7.3(36)** |  |  |  |
| **Daily use of THC=>10%** | **3.5(28)** | **20.7(117)** |  |  |  |

### Proportion of the variance explained between cases and controls by patterns of cannabis use and SZ PRS

A model including the SZ PRS, 10 PCs plus age, sex and sites explained R^2^=11.3% (R^2^- Diff: p<0.001) of the variance between cases and controls. A model including age, sex, sites, tobacco smoking and daily use of cannabis explained R^2^= 15% (R^2^- Diff: p<0.001), which was not increased by adding age at first cannabis use, R^2^=15.7% (R^2^- Diff: p=0.285). In contrast, adding data on the potency of the cannabis used to the model explained a greater proportion of the variance R^2^=17.2% (R^2^- Diff: p=0.01), which further increased to 19.2% when we first only added to the model the 10 PCs, and then to R^2^=23.0% (R^2^- Diff: p<0.001) when we also added the SZ PRS (**Figure 3**). Furthermore, ROC analyses indicated that the model only with SZ PRS, 10 PCs, age, sex and sites correctly classified 68.3% of cases, with a positive predictive value, PPV=55.2% and a negative predictive value, NPV=65.4%. This improved to 74.8% correctly classified cases, with a PPV=69.1% and NPV=75.3% for the model adding to SZ PRS, 10 PCs, age, sex, site both information on daily frequency of use and on the potency of the cannabis used.

**Figure 3:**
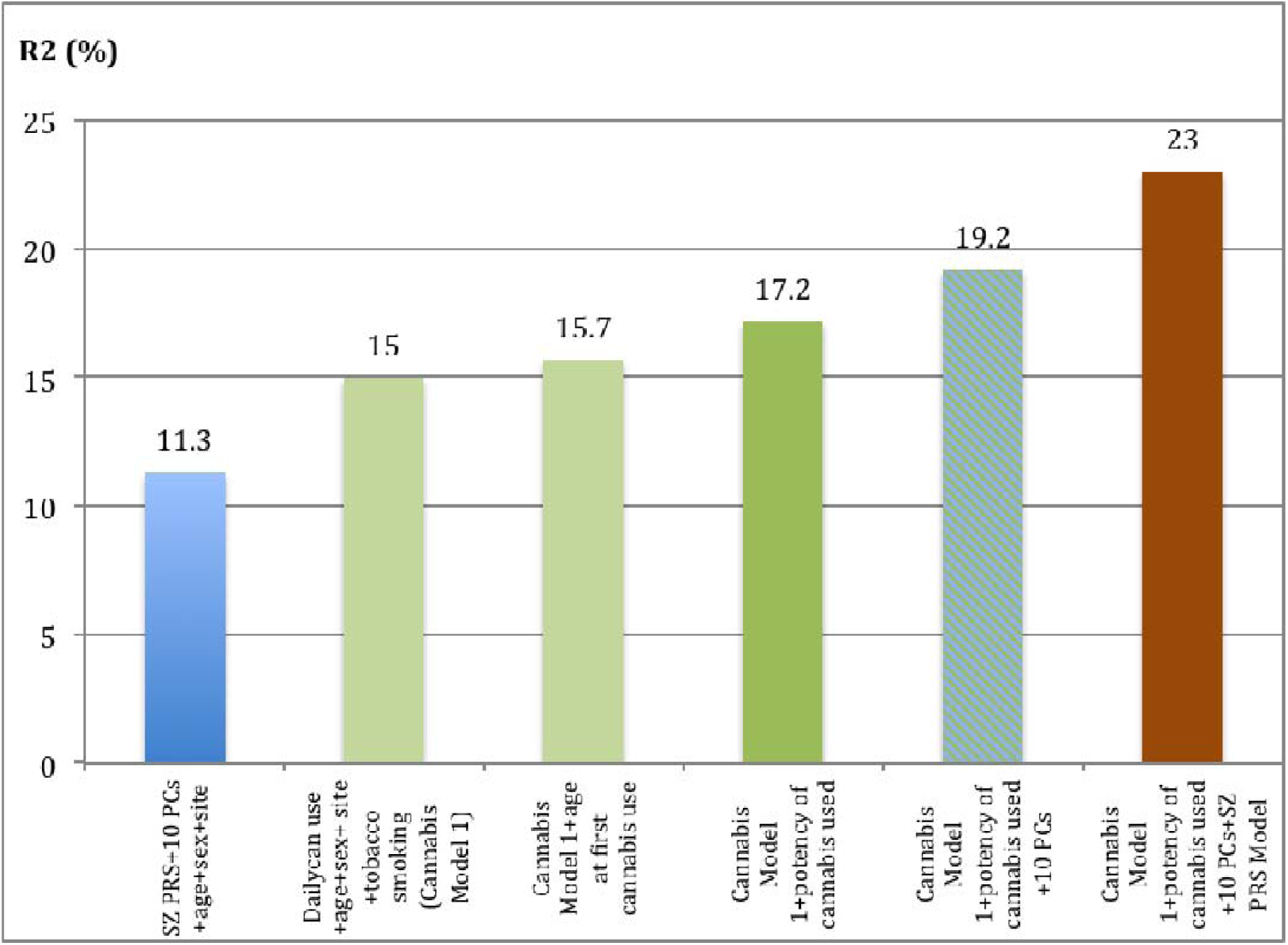
This bar chart reports the psychosis case-control variance explained (R^2^) by patterns of cannabis use and SZ PRS. The 1^st^ bar shows the R^2^ explained by the SZ PRS+ 10PCs+age+sex+site; the 2nd bar shows the R^2^ for the cannabis model 1 (Daily use+age+sex+site+tobacco smoking); the 3rd bar shows the R^2^ for the Cannabis Model 1 + age at 1st cannabis use; the 4th bar shows the R^2^ for Cannabis Model 1 + potency of cannabis use; the 5^th^ bar shows the R^2^ for Cannabis Model 1 + potency of cannabis use+ 10 PCs and the 6^th^ bar the R^2^ for the model including the Cannabis Model 1 +potency of cannabis used+ 10 PCs + SCZ PRS.

In our control sample alone, SZ PRS and 10 PCs explained a non-significant proportion of the variance between those who never used cannabis and a) having tried it at least once (lifetime use R^2^=1.9%; p=0.334), b) having started at age 15 or younger (R^2^=2.3%; p=0.557), c) having used it daily (R^2^=1.3%; p=0.678) and d) using high potency types (R^2^=2%; p=0.063).

### Does SZ PRS predict cannabis initiation and/or patterns of cannabis use?

Regression adjusted for age, sex, tobacco smoking, sites and for the 10 PCs showed that SZ PRS did not predict cannabis initiation (life time cannabis use yes/no) or starting using it at age 15 or younger both in controls (Life time cannabis use: b=0.027; p=0.51; age at 1^st^ use: b= 0.012; p=0.55) and in FEPp (Life time cannabis use b=0.001, p=0.93; age at 1^st^ use: b= −0.007;p=0.78). SZ PRS did not explain how frequently controls (b=0.027; p=0.06) or FEPp (b=0.006; p=0.91) used cannabis even when we specifically compared never use with daily use (FEPp: b=-0.013; p=0.64; Controls: b=0.003; p=0.86). Finally, SZ PRS did not predict the type of cannabis used by controls (b = 0.032; p=0.31) or by FEPp (b= 0.005; p=0.89) (supplementary Figure 1).

### The independent and combined effect of SZ PRS and pattern of cannabis use on the ORs for Psychotic Disorder

Adjusted Logistic regressions showed that SZ PRS 4^th^ *quintile* (OR=1.7; 95% CI 1.24-3.23; p=0.002) and 5^th^ *quintile* (OR=3.2; 95% CI 2.11-5.68); p<0.001) were associated with an increase in ORs for psychotic disorder compared to the 3^rd^ *quintile* (middle *quintile*) after controlling for age at 1^st^ cannabis use, frequency of use and the potency of the cannabis used. Lifetime cannabis use (crude OR=1.3, 95% CI 1.01 −1.703, p<0.050; adjusted OR=0.81, 95% CI 0.59-1.01;p=0.082) and age at 1^st^ use=<15 years old (crude OR=2.0, 95% CI 1.44-2.97, p<0.001; adjusted OR=1.1, 95% CI 0.67-1.68,p=0.771) were no longer associated with an increase in the ORs for psychotic disorder after taking into account frequency of use and cannabis potency. On the contrary, using cannabis daily and using high potency increased the ORs for psychotic disorder independently of each other, of age at 1^st^ use and also of SZ qPRS. **Table 2**

**Table 2:** This table reports the independent effect of a) SZ qPRS (quintiles) from daily cannabis use and use of high potency cannabis and b) of daily use and potency of the cannabis used from SZ PRS (quintiles) on the ORs for psychotic disorder. * *Both Crude and Adjusted ORs are controlled for: age, sex, site, smoking status and 10 PCs*

|  | Controls<br>N=787<br>(%;N) | FEP<br>N=492<br>(%;N) | *Crude-OR<br>(95% CI) | P<br>value | *Adj-OR<br>(95% CI) | P value | Interaction Statistics<br>Likelihood Ratio<br>Test (LR) | P<br>value |
| --- | --- | --- | --- | --- | --- | --- | --- | --- |
| **SZ qPRS**<br>*1<sup>st</sup> Quintile* | 24.1(190) | 13.4(66) | 3<sup>rd</sup> Quintile<br>OR=1 |  | 3<sup>rd</sup> Quintile OR=1<br>(Further adjusted<br>for frequency of<br>use, cannabis<br>potency) | 0.263 |  |  |
|  |  |  | 0.72<br>(0.49-1.06) | 0.09 | 0.74<br>(0.44-1.24) |  |  |  |
| *2<sup>nd</sup> Quintile* | 20.5(161) | 19.3(95) | 3<sup>rd</sup> Quintile OR=1 |  | 3<sup>rd</sup> Quintile OR=1 | 0.184 |  |  |
|  |  |  | 1.1<br>(0.85-1.77) | 0.266 | 1.31<br>(0.81- 2.11) |  |  |  |
| *4<sup>th</sup> Quintile* | 18.7(147) | 22.2(109) | 3<sup>rd</sup> Quintile OR=1 |  | 3<sup>rd</sup> Quintile3<br>OR=1 | 0.002 |  |  |
|  |  |  | 1.5<br>(1.19- 2.66) | 0.005 | 1.7<br>(1.24-3.23) |  |  |  |
| *5<sup>th</sup> Quintile* | 14.7(116) | 28.3(139) | 3<sup>rd</sup> Quintile OR=1 |  | 3<sup>rd</sup> Quintile OR=1 | <0.001 |  |  |
|  |  |  | 2.5<br>(1.74- 3.51) | <0.001 | 3.2<br>(2.11- 5.68) |  |  |  |
| **Daily**<br>**Cannabis**<br>**use** | 5.9 (46) | 30.0<br>(148) | Never used OR=1 |  | Never used OR=1<br>(Further adjusted<br>for SZ PRS<br>quintiles and<br>potency of use) |  | Daily use*SZ qPRS:<br>LR chi2(4) = 0.97 | 0.91 |
|  |  |  | 4.56<br>(1.59- 4.39) | <0.001 | 3.83<br>(2.97- 7.86) | <0.001 |  |  |
| **Cannabis**<br>**Potency**<br>**THC=>10**<br>**%**<br>**(High**<br>**Potency)** | 23.5<br>(185) | 45.2(223) | Never used OR=1 |  | Never used OR=1<br>(Further adjusted<br>for SZ PRS<br>quintiles and<br>frequency of use) |  | Potency*SZ qPRS:<br>LR chi2(4) = 4.98 | 0.28 |
|  |  |  | 1.72<br>(1.23- 2.29) | 0.001 | 1.8<br>(1.05- 2.71) | 0.002 |  |  |

The ORs for psychotic disorder of daily cannabis users (Interaction: Daily use*SZ qPRS: LR chi2 (4) =0.97;p=0.91) or users of high potency types (Interaction: Cannabis Potency*SZ qPRS: LR chi2(4) = 4.98;p=0.289) compared to never users were not modified by SZ PRS quintiles (qPRS). **Table 2**

Those who used potent types of cannabis more than once a week, THC=>10%, (OR=2.5,95% CI 1.26-5.00, p=0.008) or used daily either low potency, THC<10% (OR=3.5; 95% CI 1.74-7.46, p=0.001) or high potency types (OR=5.4; 95% CI 3.21-10.63,p<0.001) had an increase in the ORs for psychotic disorder compared to never users, independently of their SZ PRS and after adjusting for age, sex, site, smoking status and 10PCS. Indeed, we *plotted predictive margins* to display if and how, the probability (Pr) of being a FEPp, varied, on average, with the increase in SZ PRS across each group of the *frequency-type* composite cannabis use measure **Figure 4.** This showed that on average the probability (Pr) of being a FEPp progressively increased with the increase in SZ PRS across all the 7 groups of the *frequency-type* composite cannabis use measure. Daily users of high potency cannabis (THC=>10%) were the group with the highest Pr of being a FEPp at all level of SZ PRS, followed by daily users of THC<10% and weekly users of THC=>10%. The remaining 3 groups, never used and used rarely (i.e. less than weekly) either type of cannabis, showed a similar change in the Pr of being a FEPp with the increase in SZ PRS.

**Figure 4:**
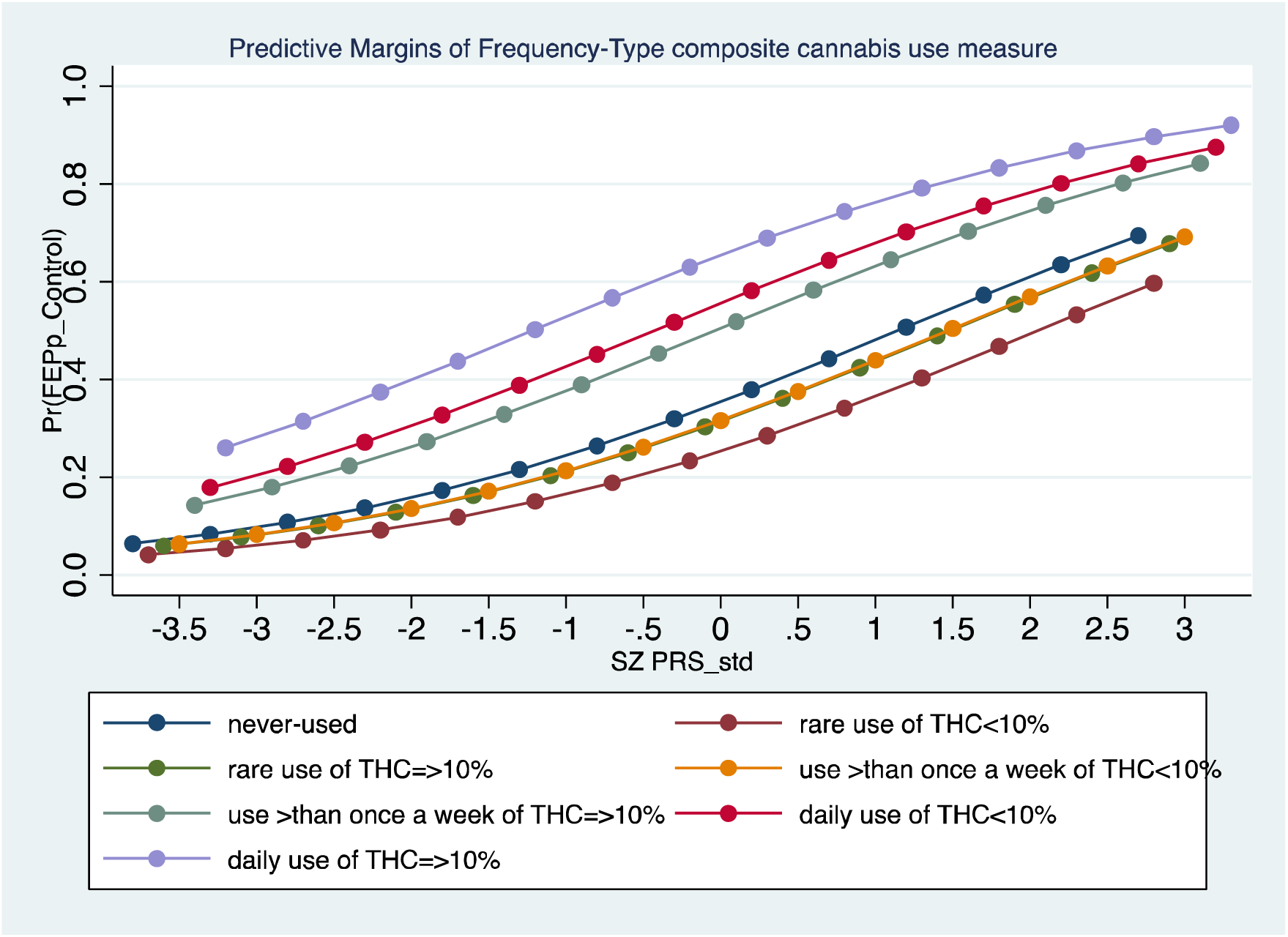
This *marginplot* describes, on average, a steady increase of the probability (Pr) of being a FEPp (y=axis) with the increase in SZ PRS (y=axis) across all the 7 groups of the frequency-type composite cannabis use measure.

Finally, an adjusted regression model with a fitted interaction term between SZ PRS and the *frequency-type* composite cannabis use measure, both fitted as continuous variables, showed an independent effect of SZ PRS (OR=2.29; 95%CI 1.71-3.05) and of the *frequency-type* composite cannabis use measure (OR=1.32; 95% CI 1.22-1.44) on the OR for psychotic disorder, but only weak evidence of an interaction (“*prs_std*\**Frequency_Type”* OR=0.91, 95% CI 0.84-0.99; p=0.033; Likelihood ratio test LR chi2(1) = 4.50; p= 0.033).

### Heavy cannabis user-only analyses

Within the sample of cannabis users only, adjusted logistic regression indicated a trend for increase in ORs for Psychotic Disorder in daily cannabis users in the 4^th^ (OR=2.6; 95% CI 0.91-9.84;p=0.06) and 5^th^ (OR=3.1; 95% CI 0.74-11.9; p=0.08) SZ PRS *quintiles* compared to those in the 3^rd^ *quintiles*. The same pattern was shown among users of high potency cannabis with those in the 4^th^ SZ PRS quintile (OR=2.7; 95%CI1.2-6.1; p=0.01) and in the 5^th^ quintile (OR=3.6; 95%CI 1.4-9.0; p=0.006), compared to the 3^rd^ one, reaching a significant increase in the ORs for Psychotic Disorder (**Table 3**).

**Table 3:**
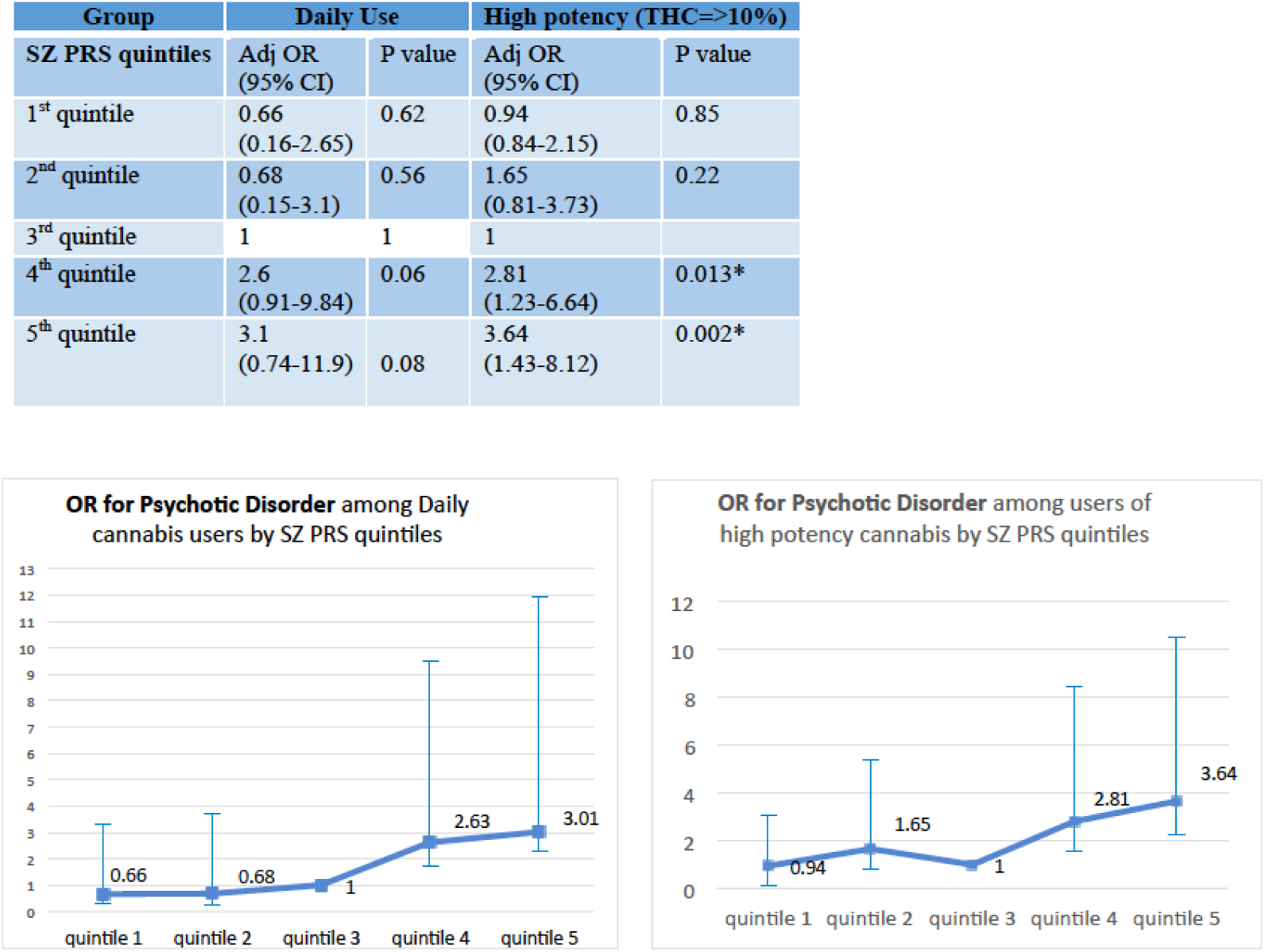
This table and corresponding graphs show the variation in the Adjusted* ORs (Y axes) for Psychotic Disorder by SZ-PRS quintiles (middle quintile as reference) for: 1) those who used cannabis daily and 2) those who used high potency types (THC=>10%). *(*Adjusted for age, gender, tobacco smoking, site and 10 PCs*)

| Group | Daily Use |  | High potency (THC=>10%) |  |
| --- | --- | --- | --- | --- |
| SZ PRS quintiles | Adj OR (95% CI) | P value | Adj OR (95% CI) | P value |
| 1<sup>st</sup> quintile | 0.66 (0.16-2.65) | 0.62 | 0.94 (0.84-2.15) | 0.85 |
| 2<sup>nd</sup> quintile | 0.68 (0.15-3.1) | 0.56 | 1.65 (0.81-3.73) | 0.22 |
| 3<sup>rd</sup> quintile | 1 | 1 | 1 |  |
| 4<sup>th</sup> quintile | 2.6 (0.91-9.84) | 0.06 | 2.81 (1.23-6.64) | 0.013* |
| 5<sup>th</sup> quintile | 3.1 (0.74-11.9) | 0.08 | 3.64 (1.43-8.12) | 0.002* |

ROC analysis indicated that among daily users of high potency cannabis (THC=>10%) a model including age, sex, site and smoking status correctly classified 83.4% of FEPp, with a Sensitivity=98.7%, a Specificity=28.6%, a PPV=84.2% and a NPV= 85.7%. When adding SZ PRS and its 10PCs to the information on age, sex, site and tobacco smoking, the proportion of correctly classified FEPp increased to 86.4% with a Sensitivity=92.9%, a Specificity=64.7%, a PPV= 89.83% and a NPV= 73.3%.

## Discussion

Our findings are the first providing estimates of risk for psychotic disorder by joint modelling the severity of cannabis use and common variant genetic liability to schizophrenia.

Indeed and in contrast with the reports from two of the MR studies ^16,17^ that suggest a causal relationship between schizophrenia genes and cannabis initiation, we found that SZ PRS did not predict individuals’ propensity to try cannabis, age at first use, frequency of use, or the potency of the cannabis used. All analyses took into account the effect of age, sex, study sites, tobacco smoking and 10PCs. These findings are consistent with a recent cross-sectional study, which tested gene-environment interaction between SZ PRS and regular cannabis use and showed no evidence of correlation between the two ^34^.

Several large genetic studies have reported that SZ PRS explained around 1% or less of the variance in life-time cannabis use and or CUD ^13,14^. In our control sample we found that SZ PRS explained a similar, but non-significant proportion of the variance for all our measures of cannabis.

As Hamilton et al. have discussed, different sample populations can present with significant differences in the variance of both the independent variable, here cannabis use and of the independent variable, SZ PRS, resulting in R^2^, which are not always comparable between samples ^35^. For instance, in our sample, the *site* where controls lived alone explained a significant 6% (p<0.001) of the variance in use of high potency cannabis, suggesting the important impact of the environmental context, such as known differences across our sites in availability of high potency cannabis ^22^, on who is likely to use it.

The evidence that the genetics of cannabis use and schizophrenia overlap does not explain why only a minority of cannabis users, even of those using daily and using high potency types ^7,21^ develop a psychotic disorder. We addressed this question by testing for interaction between SCZ PRS and measures of cannabis use, controlling for the confounding effect of study sites, age, sex, tobacco smoking and 10PCs. Firstly in our sample, SZ PRS, daily cannabis use and use of high potency independently from each other increase the ORs for psychotic disorders. Hence, daily users of high potency cannabis had an over 5-fold increase in the OR for psychotic disorder compared to never users even when controlling for SZ PRS. Furthermore, we found no evidence that SZ PRS modified the effect of lifetime cannabis use, age at 1^st^ use, frequency of use and type of potency used on the OR for psychotic disorder. Though, we report weak evidence that SZ PRS might modify the effect of the *frequency-type* composite cannabis use measure on the OR for psychotic disorder.

In **Figure 4** we report that on average the probability (Pr) of suffering from a psychotic disorder progressively increased with the increase in SZ PRS across all the seven groups of the *frequency-type* composite cannabis use measure. In line with the existing evidence^6^, those who used high potency cannabis (THC=>10%) daily were the group with the highest Pr of being a FEPp at all levels of SZ PRS, followed by daily users of THC<10% and weekly users of THC=>10%. However, we found no evidence of a positive interaction between the *frequency-type* composite cannabis use measure and the SZ PRS.

Our data indicate that SZ PRS and heavy cannabis use (e.g. daily use, use of high potency types) exert effects independent from each other on the OR for psychotic disorder. Moreover, to date the analyses testing for overlap, correlation and direction of causality between the genetics of Schizophrenia and the genetics of cannabis use have relied on data from the PGC 2 SZ GWAS. The latter is likely to have included more cannabis users among the cases than the controls, as it has been consistently reported that patients with schizophrenia have higher rates of cannabis use than the general population ^36,37^. This could partially explain the reported shared genetics and have confounded the findings suggesting a direction of causality from SZ genes to some cannabis use phenotypes.

Following from existing evidence that indicates individuals at high risk for psychotic disorder ^38^ and/or with a known family for psychosis ^39^, are more vulnerable to the psychotogenic effect of cannabis use, we show that adding SZ PRS data to easily available socio-demographic information could improve the identification of those heavy cannabis users who are more likely to suffer from a psychotic disorder. Not all heavy cannabis users develop a psychotic disorder, though we have previously shown ^5^ that daily frequency of use and use of high potency cannabis account for a significant proportion of new cases of psychotic disorder across Europe. Therefore, beginning to identify a set of data, including genetic summary scores like SZ PRS, that more accurately classify those daily uses of high potency cannabis at risk of psychotic disorder, could have important public health implications.

Our findings need to be appraised in the context of our study’s strengths and limitations. For instance the study sample size, reduced to obtain a more ethnically homogenous population, might have affected the power to detect a significant interaction between the single categorical measures of cannabis use, daily use, age at first use and use of high potency and SZ PRS.

Another limitation may be the lack of biological measures validating our self reported data of cannabis use. Nevertheless, as we were interested not on the effect of recent use but on life time exposure, measures such as urine, blood or hair samples testing could not have validated history of use over previous years ^40,41^. Furthermore, studies that have collected both laboratory and self-reported information on cannabis use have shown both sets of data to be highly correlated ^42^.

Our estimates of cannabis potency cannot account for differences in the THC% in individual samples. Our cut off of THC =10% is conservative and likely to have resulted in an underestimate of the effects of cannabis potency on the ORs for psychotic disorder.

Importantly, our study is the first to test the relationship between SZ PRS and use of high potency cannabis; the latter known to be increasing worldwide, and to be associated, with high rates of psychosis across Europe ^7^. Moreover, in our previous paper we described a probabilistic sensitivity analyses, which showed that selection bias is unlikely to explain the reported findings on the strength of the impact of daily cannabis use and use of high potency on the ORs for psychotic disorder ^7^.

Findings from first episode studies are a) less likely to be biased by illness course and less likely to produce recall bias than other study designs relying on history of exposure to environmental factors that is collected retrospectively as in prevalence samples ^43^. Finally, an important strength of our study lies in our control samples, which were recruited to represent the population at risk of each of the study sites catchment area ^7,18^. This was achieved by setting quotas based on the main socio-demographics of the populations at risk.

In conclusion, our findings suggest that despite reports of an overlap between the genetics of schizophrenia and of cannabis use, SZ PRS is far from explaining who is going to use cannabis and their pattern of use. At a time when cannabis use is increasing in popularity and becoming accessible even as a prescription drug, our study provides the first indication that using genetic data might become a tool to guide how much cannabis (and containing how much THC) an individual with a certain SZ PRS can safely use, and how likely they are to develop psychotic disorder if they use high potency cannabis daily.

## References

1. Grucza RA, Agrawal A, Krauss MJ, Cavazos-Rehg PA, Bierut LJ. Recent Trends in the Prevalence of Marijuana Use and Associated Disorders in the United StatesPrevalence of Marijuana Use and Associated Disorders in the USLetters. JAMA Psychiatry. 2016;73(3):300–301.

2. Hall W, Lynskey M. Evaluating the public health impacts of legalizing recreational cannabis use in the United States. Addiction. 2016;111(10):1764–1773.

3. Freeman TP, Groshkova T, Cunningham A, Sedefov R, Griffiths P, Lynskey MT. Increasing potency and price of cannabis in Europe, 2006–16. Addiction. 2019;114(6):1015–1023.

4. Gage SH, Hickman M, Zammit S. Association Between Cannabis and Psychosis: Epidemiologic Evidence. Biological Psychiatry. 2016;79(7):549–556.

5. Murray RM, Englund A, Abi-Dargham A, et al. Cannabis-associated psychosis: Neural substrate and clinical impact. Neuropharmacology. 2017;124:89–104.

6. Marconi A, Di Forti M, Lewis CM, Murray RM, Vassos E. Meta-analysis of the Association Between the Level of Cannabis Use and Risk of Psychosis. Schizophrenia Bulletin. 2016;42(5):1262–1269.

7. Di Forti M, Quattrone D, Freeman TP, et al. The contribution of cannabis use to variation in the incidence of psychotic disorder across Europe (EU-GEI): a multicentre case-control study. The Lancet Psychiatry. 2019;6(5):427–436.

8. Kendler KS, Ohlsson H, Sundquist J, Sundquist K. Prediction of Onset of Substance-Induced Psychotic Disorder and Its Progression to Schizophrenia in a Swedish National Sample. American Journal of Psychiatry. 2019;176(9):711–719.

9. Agrawal A, Lynskey MT. The genetic epidemiology of cannabis use, abuse and dependence. Addiction. 2006;101(6):801–812.

10. Agrawal A, Neale MC, Jacobson KC, Prescott CA, Kendler KS. Illicit drug use and abuse/dependence: modeling of two-stage variables using the CCC approach. Addictive Behaviors. 2005;30(5):1043–1048.

11. Verweij KJH, Zietsch BP, Lynskey MT, et al. Genetic and environmental influences on cannabis use initiation and problematic use: a meta-analysis of twin studies. Addiction (Abingdon, England). 2010;105(3):417–430.

12. Kendler KS, Ohlsson H, Maes HH, Sundquist K, Lichtenstein P, Sundquist J. A population-based Swedish Twin and Sibling Study of cannabis, stimulant and sedative abuse in men. Drug Alcohol Depend. 2015;149:49–54.

13. Demontis D, Rajagopal VM, Thorgeirsson TE, et al. Genome-wide association study implicates CHRNA2 in cannabis use disorder. Nature Neuroscience. 2019;22(7):1066–1074.

14. Power RA, Verweij KJH, Zuhair M, et al. Genetic predisposition to schizophrenia associated with increased use of cannabis. Molecular psychiatry. 2014;19(11):1201–1204.

15. Vaucher J, Keating BJ, Lasserre AM, et al. Cannabis use and risk of schizophrenia: a Mendelian randomization study. Molecular psychiatry. 2018;23(5):1287–1292.

16. Gage SH, Jones HJ, Burgess S, et al. Assessing causality in associations between cannabis use and schizophrenia risk: a two-sample Mendelian randomization study. Psychological medicine. 2017;47(5):971–980.

17. Pasman JA, Verweij KJH, Gerring Z, et al. GWAS of lifetime cannabis use reveals new risk loci, genetic overlap with psychiatric traits, and a causal influence of schizophrenia. Nature neuroscience. 2018;21(9):1161–1170.

18. Jongsma HE, Gayer-Anderson C, Lasalvia A, et al. Treated incidence of psychotic disorders in the multinational eu-gei study. JAMA Psychiatry. 2018;75(1):36–46.

19. McGuffin P, Farmer A, Harvey I. A polydiagnostic application of operational criteria in studies of psychotic illness: Development and reliability of the opcrit system. Archives of General Psychiatry. 1991;48(8):764–770.

20. Quattrone D, Di Forti M, Gayer-Anderson C, et al. Transdiagnostic dimensions of psychopathology at first episode psychosis: findings from the multinational EU-GEI study. Psychological Medicine. 2018:1–14.

21. Di Forti M, Marconi A, Carra E, et al. Proportion of patients in south London with first-episode psychosis attributable to use of high potency cannabis: a case-control study. The Lancet Psychiatry. 2015;2(3):233–238.

22. European Monitoring Centre for Drugs and Drug Addiction (EMCDDA). European Drug Report 2016: Trends and Developments. Publications Office of the European Union. Luxembourg 2016.

23. Niesink RJ, Rigter S. THC-concentraties in wiet, nederwiet en hasj in Nederlandse coffeeshops (2012-2013). AF1221. In. Utrecht: Trimbos-instituut; 2013.

24. Niesink RJ, Rigter S, Koeter MW, Brunt TM. Potency trends of Δ9-tetrahydrocannabinol, cannabidiol and cannabinol in cannabis in the Netherlands: 2005–15. Addiction. 2015;110(12):1941–1950.

25. Observatoire Français des Drogues et des Toxicomanies (OFDT). Drogues, chiffres clés. Paris 2015.

26. Zamengo L, Frison G, Bettin C, Sciarrone R. Cannabis potency in the Venice area (Italy): Update 2013. Drug Testing and Analysis. 2015;7(3):255–258.

27. de Oliveira GL, Voloch MH, Sztulman GB, Neto ON, Yonamine M. Cannabinoid contents in cannabis products seized in São Paulo, Brazil, 2006–2007. Forensic Toxicology. 2008;26(1):31–35.

28. Potter DJ, Clark P, Brown MB. Potency of Δ9–THC and Other Cannabinoids in Cannabis in England in 2005: Implications for Psychoactivity and Pharmacology*. Journal of Forensic Sciences. 2008;53(1):90–94.

29. Potter DJ, Hammond K, Tuffnell S, Walker C, Forti MD. Potency of Δ9– tetrahydrocannabinol and other cannabinoids in cannabis in England in 2016: Implications for public health and pharmacology. Drug Testing and Analysis. 2018;10(4):628–635.

30. Hardwick S, King S. Home Office Cannabis Potency Study 2008. London: Home Office Scientific Development Branch; 2008.

31. Schizophrenia Working Group of the Psychiatric Genomics C, Ripke S, Neale BM, et al. Biological insights from 108 schizophrenia-associated genetic loci. Nature. 2014;511:421.

32. Lewis CM, Vassos E. Prospects for using risk scores in polygenic medicine. Genome Med. 2017;9(1):96–96.

33. Vassos E, Di Forti M, Coleman J, et al. An Examination of Polygenic Score Risk Prediction in Individuals With First-Episode Psychosis. Biological Psychiatry. 2017;81(6):470–477.

34. Guloksuz S, Pries L-K, Delespaul P, et al. Examining the independent and joint effects of molecular genetic liability and environmental exposures in schizophrenia: results from the EUGEI study. World Psychiatry. 2019;18(2):173–182.

35. Hamilton DF, Ghert M, Simpson AHRW. Interpreting regression models in clinical outcome studies. Bone Joint Res. 2015;4(9):152–153.

36. Carey CE, Agrawal A, Bucholz KK, et al. Associations between Polygenic Risk for Psychiatric Disorders and Substance Involvement. Frontiers in Genetics. 2016;7(149).

37. Thornicroft G. Cannabis and Psychosis: Is there Epidemiological Evidence for an Association? British Journal of Psychiatry. 2018;157(1):25–33.

38. Vadhan NP, Corcoran CM, Bedi G, Keilp JG, Haney M. Acute effects of smoked marijuana in marijuana smokers at clinical high-risk for psychosis: A preliminary study. Psychiatry Research. 2017;257:372–374.

39. Henquet C, Murray R, Linszen D, van Os J. The Environment and Schizophrenia: The Role of Cannabis Use. Schizophrenia Bulletin. 2005;31(3):608–612.

40. Taylor M, Sullivan J, Ring SM, Macleod J, Hickman M. Assessment of rates of recanting and hair testing as a biological measure of drug use in a general population sample of young people. Addiction (Abingdon, England). 2017;112(3):477–485.

41. Curran HV, Hindocha C, Morgan CJA, Shaban N, Das RK, Freeman TP. Which biological and self-report measures of cannabis use predict cannabis dependency and acute psychotic-like effects? Psychological Medicine. 2018;49(9):1574–1580.

42. Freeman TP, Morgan CJA, Hindocha C, Schafer G, Das RK, Curran HV. Just say ‘know’: how do cannabinoid concentrations influence users’ estimates of cannabis potency and the amount they roll in joints? Addiction. 2014;109(10):1686–1694.

43. Marshall M, Lewis S, Lockwood A, Drake R, Jones P, Croudace T. Association Between Duration of Untreated Psychosis and Outcome in Cohorts of First-Episode Patients: A Systematic Review. JAMA Psychiatry. 2005;62(9):975–983.

