## Supplementary methods for "The independent and combined influence of schizophrenia polygenic risk score and heavy cannabis use on risk for psychotic disorder: A case-control analysis from the EUGEI study"

**Supplementary Material:**

**Methods**

**Recruitment:**

*Cases*: We followed procedures previously used to generate representative samples of first episode psychosis patients (FEPp) (1). We identified all individuals aged 18 to 64 years, who contacted mental health services for a suspected first episode of psychosis (FEP), over periods up to four years in 17 catchment areas in England (Southeast London, Cambridgeshire & Peterborough); France (20th arrondissement of Paris, Val-de-Marne, Puy-de-Dôme); the Netherlands (Central Amsterdam, Gouda & Voorhout); Italy (part of the Veneto region, Bologna, and Palermo); Spain (Madrid-Vallecas, Barcelona, Valencia, Oviedo, Santiago, Cuenca), and; Brazil (Ribeirão Preto, Sao Paulo) (full details of the incidence sample recruitment and general description of the incidence study methods are available from the recently published paper by Jongsma et al 2008 (2).

Case ascertainment involved trained researchers making regular contact with all secondary and tertiary mental healthcare providers to identify potential cases and searching electronic clinical records, where available. In this process, all cases with psychosis within services were considered. In all countries, it was uncommon for people to be treated for FEP in primary care; instead people with suspected psychosis would typically be referred to specialist mental health services. Research teams were overseen by a psychiatrist with experience in epidemiological research, and included trained research nurses and clinical psychologists. Teams received training in epidemiological principles and incidence study design to minimize non-differential ascertainment bias across different local and national healthcare systems (see training package on the study website: ([https://www.kcl.ac.uk/ioppn/depts/hspr/research/social-epidemiology-research-group/current-projects.aspx](https://emea01.safelinks.protection.outlook.com/?url=https%3A%2F%2Fwww.kcl.ac.uk%2Fioppn%2Fdepts%2Fhspr%2Fresearch%2Fsocial-epidemiology-research-group%2Fcurrent-projects.aspx&data=01%7C01%7Cmarta.diforti%40kcl.ac.uk%7C122825d750d549175d9908d6488b8d57%7C8370cf1416f34c16b83c724071654356%7C0&sdata=ZFnLEzDAzK8NVlXuraP4pqgV%2FKpFPwT1DVIHePwld6E%3D&reserved=0)) .

As explained in the main text, between May 1, 2010, and April, 1 2015, we approached 1519 patients with first-episode psychosis. Of these 356 (1%) refused to participate, 19 (23%) could not consent because of language barriers and 14 (0·9%) were later excluded (London N=3; Madrid N=2; Bologna N=1; Ribeirão Preto N=8) as they did not meet the age inclusion criteria. For all patients who were not part of the study, local research ethics committees approved the extraction of demographics and clinical information from patient records. Patients who refused to participate were older [FEP_consented_ mean age = 30·8(10.5), median=29(22 to 37); FEP_refused_ mean age= 32·8(11·5), median=31(25 to 42); p<0·0001], more likely to be women [FEP_consented_ male = 558(61·9%); FEP_refused_ male 311 (54·7%), χ2(1)=7·6; p=0·00603] and of White European origin [χ2(5)=38,p<0·0001] (s-Table2 for details by site).

1130 First Episode Psychosis Patients (FEPp) across the study sites consented to take part in the case-control study (s-**Table 1)**. The FEPp recruited in the case-control study are broadly representative for gender and ethnicity of the rest of the incidence sample. However, in London, Amsterdam and Ribeirao Preto cases aged 18-24 were over-represented in the case-control sample and those aged 45-54 and 55 or over were under-represented compared with the incidence sample (s-**Table 2**)

**Supplementary Table 1:** Number of participants of the case-control study recruited by each site who met the inclusion criteria.

| Catchment area |  |  |
| --- | --- | --- |
| **England** | **Controls** | **Cases** |
| Southeast London | 230 | 201 |
| Cambridgeshire | 108 | 45 |
| **The Netherlands** |  |  |
| Amsterdam | 101 | 96 |
| Gouda & Voorhout | 109 | 100 |
| **Spain** |  |  |
| Madrid | 38 | 39 |
| Barcelona | 37 | 31 |
| Valencia* | 32 | 49 |
| Oviedo* | 39 | 39 |
| Santiago* | 38 | 28 |
| Cuenca* | 38 | 18 |
| **France** |  |  |
| Paris (Maison-Blanche)* | 0 | 36 |
| Paris (Val-de-Marne) | 100 | 54 |
| Puy-de-Dome | 47 | 15 |
| **Italy** |  |  |
| Bologna | 65 | 70 |
| Verona* | 115 | 59 |
| Palermo | 100 | 58 |
| **Brazil** |  |  |
| Ribeiãro Preto | 302 | 192 |
| **Total** | **1,499** | **1,130** |

*Sites excluded for the case-control analysis because of missing data =>10%. Mason-Blanche was excluded from the case-control analysis, as they did not recruit any controls.

**Supplementary Table 2** χ^2^ and p-values for comparisons between those cases who participated in the case-control arm of the study and those who did not. The table shows how the case-control study cases are representative of the rest of the incidence sample by site. (*Age range groups included the following categories: 18-24; 25-34; 35-44; 45-54; 55-64*) (modified from 3)

|  |  | Age | | | | | Sex | | | | | | Minority status | | |
| --- | --- | --- | --- | --- | --- | --- | --- | --- | --- | --- | --- | --- | --- | --- | --- |
|  | Mean,sd; (Median)  case-control sample | Mean;  (Median)  rest of the incidence sample | χ^2^  (based on age groups) | | p-value | Male %; N  case-control sample | Male %; N  rest of the incidence sample | | χ^2^ | p-value | %; N minority  Case control | %: N minority  Rest of the incidence sample | | χ^2^ | p-value |
| England |  |  |  |  | |  |  |  | |  |  |  | |  |  |
| Southeast London | 29·6,9·4 (27) | 34·6,11·2 (33) | **31·4** | **<0·01** | | 63·2 (127) | 51·4 (112) | **5·9** | | **0·02** | 70·6 (142) | 77·1 (168) | | 2·2 | 0·13 |
| Cambridgeshire | 28·1,7·9 (26) | 32·5,12·3 (29) | 6·8 | 0·15 | | 55·6 (25) | 57·0 (126) | 0·0 | | 0·86 | 35·6 (16) | 41·8 (87) | | 0·6 | 0·44 |
| The Netherlands |  |  |  |  | |  |  |  | |  |  |  | |  |  |
| Amsterdam | 27·6,8·1 (25) | 38·2,12·5 (36) | **50·5** | **<0·01** | | 74·0 (71) | 59·9 (118) | 5·6 | | 0·18 | 70·8 (68) | 73·6 (134) | | 0·2 | 0·62 |
| Gouda & Voorhout | 31.7,11·1 (29) | 32.5,12·0 (30) | 1 | 0·9 | | 65·0 (65) | 54·6 (36) | 1·8 | | 0·18 | 17 (17) | 35·4 (23) | | **7·2** | **0·01** |
| Spain |  |  |  |  | |  |  |  | |  |  |  | |  |  |
| Madrid | 33·1,11·1 (33) | 33·9,9·6 (30) | 2·5 | 0·64 | | 69·2 (27) | 63·3 (31) | 0·3 | | 0.56 | 10·3 (4) | 12·5 (2) | | 0·1 | 0·8 |
| Barcelona | 29·4,11·3 (30) | 30·7,13·4 (28) | 2·5 | 0·63 | | 74·2 (23) | 50·7 (39) | **5** | | **0·02** | 20 (6) | 22·4 (15) | | 0·1 | 0·79 |
| Valencia | 31·5,11·4 (27) | 35·6,10·3 (35·5) | 3·3 | 0·51 | | 61·2 (30) | 20·0 (2) | **5·7** | | **0·02** | 16·3 (8) | 22·2 (2) | | 0·2 | 0·67 |
| Oviedo | 34·7,10·8 (35) | 36·0 9·7 (33) | 3·4 | 0·49 | | 51·3 (20) | 46·5 (20) | 0·2 | | 0·67 | 20·5 (8) | 12·5 (4) | | 0·8 | 0·37 |
| Santiago | 32·1,11·2 (31) | 42·9,10·4 (44) | 8·7 | 0·07 | | 64·3 (18) | 37·5 (3) | 1·8 | | 0·17 | 0 (0) | 0 (0) | | n/a | n/a |
| Cuenca | 29·2,9·5 (27) | 28·3,11·2 (25) | 0·7 | 0·88 | | 77·8 (14) | 77·8 (7) | 0·0 | | 1·00 | 16·7 (3) | 33·3% (3) | | 1 | 0·33 |
| France |  |  |  |  | |  |  |  | |  |  |  | |  |  |
| Paris (Maison-Blanche) | 31·4,10·2 (30) | 34·1,12·1 (31) | 2·9 | 0·56 | | 66·7 (24) | 70·2 (59) | 0·1 | | 0·69 | 58·3 (21) | n/a | | n/a | n/a |
| Paris (Val-de-Marne) | 31·3,10·1 (27) | 33.6, 11·2  (30) | 4·6 | 0·33 | | 61·1 (33) | 48·1 (75) | 2·7 | | 0·1 | 22·2 (12) | n/a | | n/a | n/a |
| Puy-de-Dome | 37·3,13·4 (32) | 33·7,12·7 (34) | 8·8 | 0·07 | | 60·1 (9) | 70·4 (19) | 0·5 | | 0·49 | 20·0 (3) | n/a | | n/a | n/a |
| Italy |  |  |  |  | |  |  |  | |  |  |  | |  |  |
| Bologna | 32·5,9·9 (33) | 33·3,10·5 (30) | 7·2 | 0·13 | | 50·0 (35) | 53·7 (51) | 0·2 | | 0·64 | 28·6 (20) | 29·5 (28) | | 0·0 | 0·9 |
| Veneto | 36·5,10·1 (37) | 36·6,12·3 (36·5) | 6·9 | 0·14 | | 55·9 (33) | 52·0 (26) | 0·2 | | 0·68 | 16·7 (9) | 20 (10) | | 0·2 | 0·66 |
| Palermo | 30·1,8·9 (28) | 34·5,10·2 (31) | 12·7 | 0·01 | | 58·6 (34) | 54·6 (66) | 0·3 | | 0·6 | 6·9 (4) | 14·1 (17) | | 1·9 | 0·16 |
| Brazil |  |  |  |  | |  |  |  | |  |  |  | |  |  |
| Ribeirão Preto | 32·3,11·2 (30) | 35·9,10·6 (35) | **24·1** | **<0·01** | | 56·8 (109) | 49·1 (161) | 2·9 | | 0·09 | 49·5 (95) | 33·7 (90) | | **11·5** | **<0·01** |

*Controls:* All sites contributed to the recruitment of 1499 population controls except for Maison Blanche, which consequently was excluded from the case-control analysis (s-**Table1**). Controls were recruited using a mix of random and quota sampling that aimed to obtain samples representative for age, gender and ethnicity of each site population at risk. Nevertheless, controls aged 18-34 were over-sampled and those aged 35 and over were under-sampled (χ^2^: 212·4, p<0.01, s-**Table 3**). Differences by gender and ethnicity are also reported in s-**Table 3**. As reported in the main methods section we used inverse probability weights to account for any over and under sampling of controls relative to the populations at risk; we gave each control’s data a weight inversely proportional to their probability of selection, on key demographics (age, gender, ethnicity, using census data on relevant populations). The weights were applied in all analyses.

**Supplementary** **Table 3:** Representativeness of the control sample compared with the population-at-risk *(This does not include Paris- Maison Blanche where no controls were recruited)(2*)

|  | Population at-risk | | Controls | | | |
| --- | --- | --- | --- | --- | --- | --- |
|  | *n* | Percentage | *N* | Percentage | χ^2^ | p-value |
| **Age**  18-24  25-34  35-44  45-54  55-64 | 1,828,075  3,057,640  3,058,837  2,856,614  2,152,499 | **14.1**  23.6  23.7  21.9  16.6 | 323  511  323  253  172 | **21.7**  34.3  15.6  17.0  11.5 | 212.4 | <0.01 |
| **Sex**  Male  Female | 6,337,783  6,464,653 | **49.5**  50.5 | 672  788 | **46.0**  54.0 | 7.1 | <0.01 |
| **Minority**  **status**  Majority  Minority | 9,881,660  2,917,823 | **77.2**  22.8 | 1,072  414 | **72.1**  27.9 | 21.7 | <0.01 |

*Final FEPp and Controls sample size:* FEPp N=492 and Controls N=787 of European Ancestry were included in the study. (see flow chart below).

**FEPp recruitment flow chart**:

FEPp approached N=1519

N= 33 Excluded because:

1. Language barrier N=19
2. Outside the age inclusion criteria N=14

Refused N=356

Controls recruited N=1499

FEP Recruited (consented) N=1130

N=2190 (83% of the Total) participants gave DNA samples

GWAS 98% call rate:

FEPp N=999	Controls N=1147

Ethnic groups excluded:

African Ancestry N=402

Others N=105

Final sample

FEPp N=492	Controls N=787

**Measures:**

The Cannabis Experienced Questionnaire firstly described by Barkus et al 2006 (5), was later modified (CEQmv) (6) to expand 1) questions on the pattern of use including the assessment of the type of cannabis, 2) the section on other drug use and 3) to reduce the section on the experiences following a factor analysis (6). For the EUGEI study we further modified it (CEQ_EUGEI_) to 1) include questions to assess dependence for cannabis use and other drugs, and 2) to describe use and changes in cannabis use over 3 age periods: 0-11 years old; 12-17 years old and 18 and older.

The Cannabis Experience Questionnaire (CEQ) ‘s questions we selected to construct our measures of cannabis exposure aimed to ascertain the pattern of use that described the “ most” each participant used over the period they used; thus these were mostly questions covering life-time use rather than current use. : 1) lifetime cannabis use: have you ever used cannabis yes/no; 2) age at first use of cannabis in years that in accordance with the existing literature (7) is dichotomized as in **Supplementary-Table 4** ; 4) frequency of use. 5) What type of cannabis did you mostly use? (name given in native language; see next paragraph for more details.

These were also the measures that, in an independent sample, we reported to carry the highest OR for psychotic disorder (8).

**Supplementary-Table 4:** Single measures of cannabis use included in the analyses

| Lifetime cannabis use | 0=never used | 1=Yes |  |
| --- | --- | --- | --- |
| Age at 1^st^ use of cannabis | 0=never used | 1= started at age 16years or older | 2=started at age 15 years or younger |
| Lifetime frequency of use | 0=used never or occasionally (less than once a week) | 1=used more than once a week (but less than daily) | 2=used daily |
| Type of cannabis used | 0= never used | 1= used types with THC<10% | 2=used types with THC=>10% |

**The cannabis potency variable:**

The potency variable was created using a cut off of THC=10% based on the mean THC concentration expected in the different types of cannabis available across the side sites, as reported in the EMCDDA and by the National data on cannabis potency quoted (9). Participants were asked to name in their own language the name of the type of cannabis they mostly used during their period of use.

The low-potency cannabis category (THC<10%) included hash/resin from UK and Italy, imported herbal cannabis from UK, Italy, Spain and France, Brazilian marijuana and hash and the Dutch Geimporteerde Wiet. The high-potency category (THC=>10%) included all the other types reported by the study participants in their original language street names such as: UK home-grown skunk/sensimilla UK Super Skunk, Italian home-grown skunk/sensimilla , Italian Super Skunk, the Dutch Nederwiet, Nederhasj and geimporteerde hasj, the Spanish and French Hashish (from Morocco), Spanish home-grown sensimilla, French home-grown skunk/sensimilla/super-skunk and Brazilian skunk (10-17 )

**Statistical analysis:**

*Confounder selection*: The socio-demographic variables available and in line with the existing literature^1^ were associated with case-control status.

To estimate the possible confounding effect of tobacco smoking in our analysis, we used the data on number of cigarettes smoked over the past 12 months. As for the method used to group the raw measures of cannabis exposure, we applied a logistic regression adjusted for age, gender and ethnicity, testing for an association between the raw variable on number of cigarettes smoked per day over the previous 12 months (0=never smoked; 1=smoked less than cigarettes per day; 2= smoked 10 or more cigarettes) and case-control status. Smoking less than 10 cigarettes per day was not associated with an increase in the ORs for psychotic disorder (OR=0·9; 95% CI 0·9 to 2·8) compared to never smoked, contrary to smoking 10 cigarettes or more (OR= 2·5 95% CI 1·7 to 4·2). Therefore, the variable on tobacco use entered in the main analysis model is the one described in the main methods paragraph.

We did not include alcohol use as it was not associated with case-control status as reported in our previous publication (5).

*Genotyping:* The samples were genotyped at the MRC Centre for Neuropsychiatric Genetics and Genomics in Cardiff (UK) using a custom Illumina HumanCoreExome-24 BeadChip genotyping array covering 570,038 genetic variants. We selected common SNPs with minor allele frequency >= 5%, Hardy-Weinberg Equilibrium test p-value >= 1x10^-8^ and genotyping rate of >= 99%. Subjects with genotyping rate <= 90% and mismatched sex information were discarded. Relatedness testing was performed by PLINK. Any two DNA samples with genome identity (PI-HAT > 0.1) were defined as ‘closely related’. From each identical or related pair, only the individual with higher call rates was kept.

**Results**

Here we report the regression plots adjusted for age gender and 10PCs of SZ PRS scores in controls by a) lifetime cannabis use status; b) by daily cannabis use; c) by the use of high potency; d) by the age at first use =<15 years old. These regression plots show that, in the control sample SZ PRS does not predict cannabis initiation and/or patterns of cannabis use. **Supplementary -Figure 1.**

**Supplementary Figure 1** : Regression plots *adjusted for age gender and 10PCs* of SZ PRS scores in controls by a) lifetime cannabis use status; b) by daily cannabis use; c) by the use of high potency; d) by the age at first use =<15 years old.

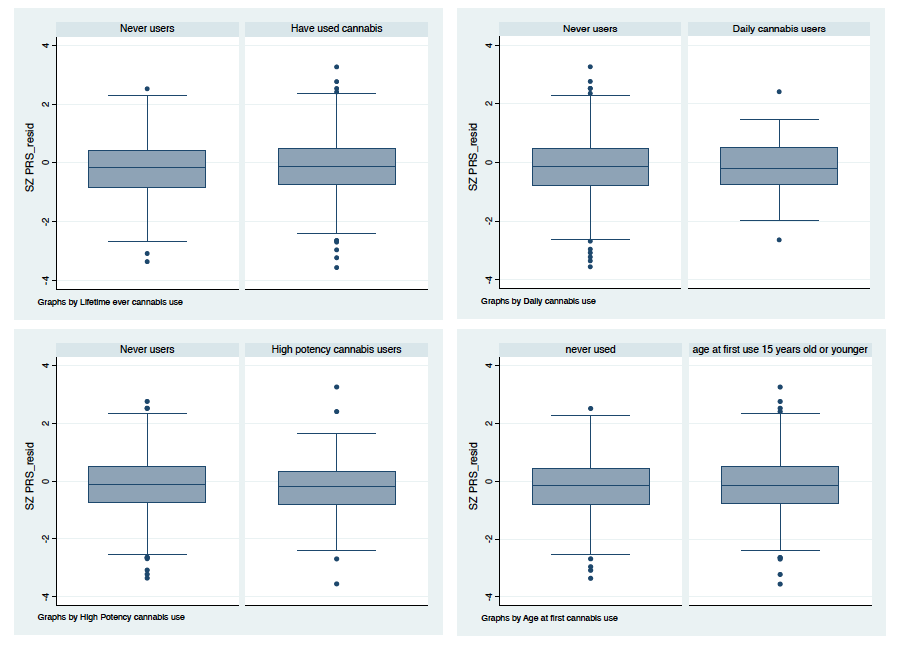

**Supplementary methods reference:**

1.	Fearon P, Kirkbride JB, Morgan C, Dazzan P, Morgan K, Lloyd T, et al. Incidence of schizophrenia and other psychoses in ethnic minority groups: results from the MRC AESOP Study. Psychological Medicine. 2006;36(11):1541-50.

2.	Jongsma HE, Gayer-Anderson C, Lasalvia A, et al. Treated incidence of psychotic disorders in the multinational eu-gei study. JAMA Psychiatry. 2018;75(1):36-46.

3.	Jongsma HE. The role of the sociocultural context in explaining variance in incidence of psychosis and higher rates of disorder in minorities. [PhD Thesis]: University of Cambridge; 2018.

4.	Vandenbroucke JP, Pearce N. Case–control studies: basic concepts. International Journal of Epidemiology. 2012;41(5):1480-9.

5.	Barkus EJ, Stirling J, Hopkins RS, Lewis S. Cannabis-induced psychosis-like experiences are associated with high schizotypy. Psychopathology. 2006;39(4):175-8.

6.	Di Forti M, Morgan C, Dazzan P, Pariante C, Mondelli V, Marques TR, et al. High-potency cannabis and the risk of psychosis. British Journal of Psychiatry. 2009;195(6):488-91.

7.	Casadio P, Fernandes C, Murray RM, Di Forti M. Cannabis use in young people: The risk for schizophrenia. Neuroscience & Biobehavioral Reviews. 2011;35(8):1779-87.

8.	Di Forti M, Marconi A, Carra E, et al. Proportion of patients in south London with first-

episode psychosis attributable to use of high potency cannabis: a case-control study. *The Lancet*

*Psychiatry.* 2015;2(3):233-238.

9. European Monitoring Centre for Drugs and Drug Addiction (EMCDDA). European Drug Report 2016: Trends and Developments. Publications Office of the European Union. Luxembourg 2016.

10.	European Monitoring Centre for Drugs and Drug Addiction (EMCDDA), Ministry of Health and Consumer Affairs (Spain). Spain National Report to the EMCDDA 2012.

11.	Niesink R RS. THC-concentraties in wiet, nederwiet en hasj in Nederlandse coffeeshops (2012-2013). AF1221. Utrecht: Trimbos-instituut; 2013.

12.	Observatoire Français des Drogues et des Toxicomanies (OFDT). Drogues, chiffres clés. Paris 2015.

13.	Zamengo L, Frison G, Bettin C, Sciarrone R. Cannabis potency in the Venice area (Italy): Update 2013. Drug Testing and Analysis. 2015;7(3):255-8.

14.	Niesink RJM, Rigter S, Koeter MW, Brunt TM. Potency trends of Δ9‐tetrahydrocannabinol, cannabidiol and cannabinol in cannabis in the Netherlands: 2005–15. Addiction. 2015;110(12):1941-50.

15.	de Oliveira GL, Voloch MH, Sztulman GB, Neto ON, Yonamine M. Cannabinoid contents in cannabis products seized in São Paulo, Brazil, 2006–2007. Forensic Toxicology. 2008;26(1):31-5.

16.	Potter DJ, Clark P, Brown MB. Potency of Δ9–THC and Other Cannabinoids in Cannabis in England in 2005: Implications for Psychoactivity and Pharmacology*. Journal of Forensic Sciences. 2008;53(1):90-4.

17.	Hardwick S KS. Home Office Cannabis Potency Study 2008. London: Home Office Scientific Development Branch; 2008.
